# Multi-parametric ex vivo magnetic resonance imaging for myelination in the developing mouse brain

**DOI:** 10.64898/2026.09.26.754665

**Authors:** Choong H. Lee, Jennifer A. Minteer, Zifei Liang, Seon-Hi Shin, Yongsoo Kim, Jiangyang Zhang

## Abstract

Existing MRI-based markers have limited sensitivity and specificity for assessing myelin in the brain. This study investigated whether multi-parametric MRI could improve myelin mapping in the developing mouse brain. T_1_, T_2_, magnetization transfer, and diffusion MRI data were acquired at 7 Tesla from ex vivo myelin-associated oligodendrocyte basic protein (MOBP)-enhanced green fluorescent protein (eGFP) mouse brains at postnatal day 14 (P14), P35, and P56. Serial two-photon tomography data from the same mouse brains were used to quantify myelin content based on the MOBP signal intensity. Although most MR parameters demonstrated significant age-related changes, only a subset correlated strongly with MOBP signals. Partial least squares regression analysis of MRI and MOBP data from the corpus callosum and other brain regions demonstrated that the multi-parametric approach improved myelin mapping in the developing mouse brain. T_2_ was ranked as the highest contributor to myelin estimation across multiple brain regions.

## 1. Introduction

Myelin ensheathes axons of the central nervous system (CNS) and facilitates fast conduction of action potential (1). Myelin consists primarily of lipids, which occupy 70-80% of the dry weight of myelin (2), and proteins, which are crucial for structural stability of the myelin (3, 4). Disruption of myelin integrity is a hallmark of many neurological diseases, occurring either as a primary target in conditions such as multiple sclerosis (MS) (5) and leukodystrophies (6, 7), or as a secondary consequence in diseases like Alzheimer’s disease (8, 9). Given myelin’s critical role in brain function and its involvement in neurological disorders, accurate assessment of myelin integrity is essential for understanding disease mechanisms and monitoring therapeutic interventions.

MRI is the preferred method for non-invasive myelin mapping because it provides multiple parameter maps that are sensitive to myelin and myelin-related injuries. These parameters include relaxivity (e.g., T_1_, T_2_) (10–15), magnetic susceptibility (16–18), magnetization transfer (MT) (19–23), and diffusion MRI metrics (24–26). More advanced myelin markers have also been developed, including those based on microstructural models such as myelin water imaging (12, 27, 28) macromolecule pool fraction (29–31), and white matter tract integrity measures (32–34), as well as advanced contrast mechanisms such as inhomogeneous magnetization transfer (ihMT) (35–37) and ultrashort echo time imaging (38–40). However, recent studies have demonstrated that individual MRI markers have limited specificity for myelin mapping (41–43), mainly due to the fact that MRI is an indirect measurement and its signals reflect multiple aspects of tissue microstructure other than myelin.

Given the limitations of single-parameter MRI approaches, multi-parametric MRI (mpMRI) has emerged as a promising solution (44). This technique aims to improve myelin specificity by combining multiple MRI contrasts (42, 44). For example, Warntjes et al. estimated myelin content based on relaxivity and proton density data (45), and Merkler et al. demonstrated that combining relaxivity and magnetization transfer better distinguishes myelin pathology in the mouse cuprizone model(46). Extending this multiparametric approach, we recently incorporated relaxivity, magnetization transfer, and diffusion metrics to study enhanced myelination in a mouse model (47–49), demonstrating improved myelin specificity (47).

Further development of mpMRI requires robust histological validation. Conventional histological processing is labor-intensive, time-consuming, and error-prone, resulting in limited availability of high-quality validation data. Due to this scarcity, previous studies correlating MRI with histology have often pooled results from multiple brain structures. Recent evidence suggests that microstructural differences among tissues may confound such analyses and lead to overestimation of correlations (41, 43). Therefore, obtaining sufficient histological data to enable structure-specific correlation analyses is essential for minimizing potential confounding factors and improving the accuracy of MRI-histology relationships.

In this study, we investigated age-related changes in MR parameters using myelin-associated oligodendrocyte basic protein (MOBP)-enhanced green fluorescent protein (eGFP) transgenic mice. MOBP is the third most abundant protein in CNS myelin and is expressed exclusively in oligodendrocytes, making the co-expressed eGFP a highly specific and quantifiable marker of myelin content that can be directly correlated with MRI parameters. We examined the relationship between MRI parameters and myelin content, measured using serial two-photon tomography (STPT)(50), at three postnatal developmental stages: P14, P35, and P56, corresponding to early, mid, and late phases of myelination.

## 2. Methods

### 2.1. Animals

All animal procedures were conducted in accordance with protocols approved by the Institutional Animal Care and Use Committee (IACUC) at Pennsylvania State University. MOBP-eGFP mice were originally generated by the GENSAT project and obtained from the Mutant Mouse Resource and Research Center (MMRRC; stock number 030483-UCD). The mouse line was generously provided to our laboratory by Dr. Shin Kang (Temple University) under an approved material transfer agreement with MMRRC. The imported MOBP-eGFP mice were crossed to C57bl/6J (JAX Strain #:000664) more than 5 generations before the experimental usage. Both male and female mice were used and housed under a 12-hour light/dark cycle at 22–25 °C with ad libitum access to food and water. MOBP-eGFP expression was confirmed using PCR of genomic tail biopsy DNA. Samples were collected at postnatal days 14, 35, and 56 (n = 5 in each age group). Mice were deeply anesthetized with ketamine (100 mg/kg) and xylazine (10 mg/kg) and transcardially perfused with 0.9% saline followed by 4% paraformaldehyde (PFA) in 0.1 M phosphate buffer (PB, pH 7.4). Decapitated heads were post-fixed in 4% PFA for 24 h at 4 °C and transferred to 0.05M PB prior to shipment for ex vivo, in-skull MRI acquisition at New York University.

### 2.2. MRI Acquisition

MRI data were acquired on a 7 Tesla MRI system equipped with a quadrature transmit volume coil (70 mm diameter) and a 4-channel receive-only phased array cryogenic probe (Bruker Biospin, Billerica, MA, USA). Mouse brains were placed in 10 ml syringes filled with Fomblin (perfluoropolyether; Ausimont USA Inc.), which has no MRI visible proton, to match tissue susceptibility and prevent dehydration. During MRI, the specimens were kept at 36–36.5 °C using a heater inside the cryogenic probe. Co-registered two-dimensional multi-parametric MRI data were acquired with an in-plane resolution of 0.1 mm x 0.1 mm, a slice thickness of 0.7 mm, and 14 slices in total.

For T_1_ and T_2_ MRI, we used the RAREVTR (Rapid Acquisition with Relaxation Enhancement and Variable Repetition time (TR)) (51) and MSME (Multi-Slice Multi-Echo) sequence with the parameters in **Table 1**. For MT-MRI, we used an offset frequency of 5 kHz for conventional MT-MRI and 10 kHz for ihMT-MRI following previous reports by Prevost et al. (7). Co-registered diffusion tensor data were acquired using a diffusion-weighted echo-planar imaging (DW-EPI) sequence. T_1_/T_2_, MT/ihMT, DW-EPI images were acquired using the scan parameters shown in **Table 1**. The ihMT parameters were chosen based on previous studies on ex vivo mouse brain under the same condition (37, 52–54).

**Table 1:** MR imaging parameters used in this study. Abbreviations are: AT: acquisition time; NEX: number of excitations; TE: echo time; TR: repetition time.

| A | TE (ms) |  | TR (ms) |  | Resolution (mm<sup>3</sup>) |  | NEX/AT |
| --- | --- | --- | --- | --- | --- | --- | --- |
| T1-weighted RARE images | 6.3 |  | 868, 1096, 1365, 1694, 2115, 2702, 3682, 7500 |  | 0.1 x 0.1 x 0.7 |  | 2/15 min |
| T2-weighted MSME images | 7, 14, 21, 28, 34, 41, 48, 55, 62, 69, 76, 83, 89, 96, 103, 110, 117, 124, 131, 138, 144, 151, 158, 165, 172 |  | 4000 |  | 0.1 x 0.1 x 0.7 |  | 2/40 min |

| B | TE (ms) | TR (ms) | Offset Freq (kHz) | Echo Train Length | Saturation Pulse | Pulsewidth/Bandwidth/Repetition time/# of pulses | Resolution (mm<sup>3</sup>) | NEX/AT |
| --- | --- | --- | --- | --- | --- | --- | --- | --- |
| MT | 5.5 | 4000 | 5 | 6 | Gaussian | 1.83 ms/1.5 kHz/2.62ms/200 | 0.1 x 0.1 x 0.7 mm | 4/22 min |
| ihMT | 5.5 | 4000 | 10 | 6 | Gaussian | 1.83 ms/1.5 kHz/2.62ms/200 | 0.1 x 0.1 x 0.7 mm | 4/1hr 14 min |

| C | TE (ms) | TR (ms) | $\Delta/\delta$ (ms) | A0 images | Directions | B values (ss/mm<sup>2</sup>) | Resolution (mm<sup>3</sup>) | NEX/AT |
| --- | --- | --- | --- | --- | --- | --- | --- | --- |
| Diffusion weighted EPI | 33 | 5000 | 7/15 | 5 | 30 | 1000, 2500, 4000 | 0.1 x 0.1 x 0.7 | 2/15 min |

### Image Analysis

MR images were reconstructed from raw data on the scanner console with trajectory correction (Paravision 6.0.1, Bruker Biospin, Billerica, MA, USA). T_1_ and T_2_ values were computed using the Image Sequence Analysis (ISA) Tool offered by Bruker Paravision. Images with no saturation pulse (M0) were acquired with images with positive/negative or dual-frequency saturation pulses ( , and ). MTR was calculated as: , and ihMTR was calculated as: . From the DWIs, diffusion tensors and diffusion kurtosis tensors were obtained per voxel (55) using the standard kurtosis-fitting approach implemented in the Diffusion Kurtosis Imaging Matlab Toolbox (https://cai2r.net/resources/software/diffusion-kurtosis-imaging-matlab-toolbox) (56) and the following parameters were derived: MD, RD, FA, mean and radial kurtoses (MK/RK). The FA image was used for selecting the region of interest (ROI) manually, including the genu and splenium of corpus callosum (gcc and scc), external capsule (ec), cerebral peduncle (cp), and motor cortex (mCX) using ROIeditor (http://www.mristudio.org). Representative images with ROIs used in this study were shown in the supplementary materials **Fig. S1**.

### 2.3. Serial two-photon tomography for histological validation

Histological validation of myelin content was achieved using serial two-photon tomography (STPT) (50, 57). Brain tissue was extracted from the skull and dissected following ex vivo MR imaging. Samples were embedded in 4% oxidized agarose and chemically stabilized to maintain structural integrity during serial vibratome sectioning and two-photon image acquisition. For tissue cross-linking, samples were incubated overnight at 4 °C in a sodium borohydride (SBH) solution consisting of 0.5-1% sodium borohydrate (NaBH_4_) in 0.05M sodium borate buffer (pH 9.0-9.5). A second 4% oxidized agarose embedding method was also used, replacing SBH with an overnight acrylamide incubation at 4°C followed by heat-activated polymerization (detailed protocol available at (58)).

Whole-brain STPT imaging was performed using a TissueCyte 1000 system (TissueVision) equipped with a two-photon laser (Chameleon Ultra II, Coherent) operating at 910 nm excitation. Fluorescence signals were collected simultaneously using a 560 nm dichroic mirror. Image stacks were acquired at 1×1 µm lateral resolution with 50 µm thick serial sectioning. Samples exhibiting substantial tissue damage or imaging artifacts were excluded from further analysis. To enable anatomical correspondence across specimens and comparison with MRI data, STPT image volumes were spatially registered to age-matched developmental brain templates. Atlas registration and anatomical alignment followed previously published STPT pipelines, including template-based registration and transformation of atlas annotations into sample space, as described in detail by Liwang et al.(57). This registration framework enables region-wise mapping of MOBP-eGFP fluorescence to standardize anatomical reference spaces derived from the Allen Institute–derived developmental brain atlas.

### 2.4. Statistical Analysis

All statistical tests were performed with Prism (GraphPad). Statistical significance was determined using the student’s *t*-test with the threshold set at 0.05. Pearson correlation coefficients and p values were calculated between MR parameters and MOBP signal intensities. Corrections for multiple comparisons were performed using the Bonferroni method. Conventional multivariable linear regression is unsuitable for identifying the relationship between MR parameters and MOBP signals due to collinearity among the MR parameters. As an alternative, we employed partial least squares regression (PLSR), which effectively handles collinearity and generates a linear estimator for myelin from the input MR parameters. The function projected the independent variables (MR parameters) and response variables (MOBP signals) into a latent space, in which linear least square regression was carried out. Latent components in PLSR are synthetic variables that condense correlated predictors into a smaller set of independent factors, each chosen to maximize its ability to explain variance in both the predictors and the responses (59). This approach reduces collinearity within the independent variables and their dimensions without reducing the prediction power (60). PLSR also offers two important metrics: the variable importance in projection (VIP) score, which identifies the MR parameters that most significantly contribute to myelin estimation, and the percentage of variance (PCTVAR) in MOBP signals explained by the latent components of the PLSR model. We cross-checked a function *plsregress* in Matlab® 9.14.0 (R2023a) with SAS programs (version 9.4) to compute the VIP score, regression coefficients, and PCTVAR.

## 3. Results

### 3.1 Multi-parametric MRI and histology of MOBP-eGFP mice

In the STPT images (**Fig. 1**), the corpus callosum (indicated by the yellow arrows) showed increased size and intensity from P14 to P56. In the MRI data (**Fig. 1**), both T_1_ and T_2_ values were higher at P14 than at P56, with the contrast between the corpus callosum and cortex inverted in the T_2_ map between P14 and P56. We also found substantial increased contrasts in MTR and ihMTR over the same period, and the diffusion MRI parameters (FA, MD, MK) showed similar trends.

**Figure 1:**
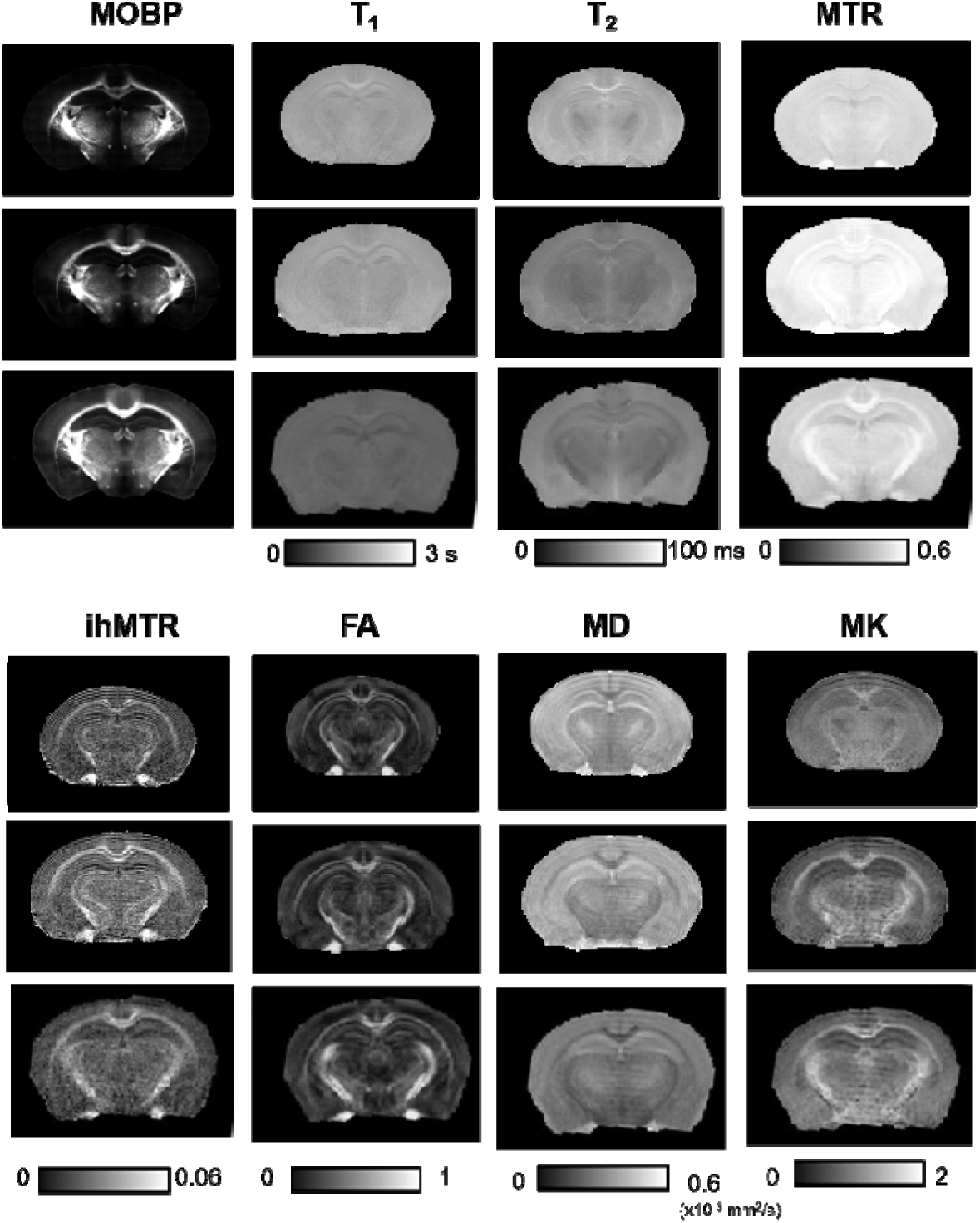
Representative STPT-MOBP images and MRI parameter maps from the MOBP-eGFP mouse brains at P14, P35, and P56.

In the gcc, MOBP signal intensity increased significantly with age (**Fig. 2**), with significant differences observed both between P14 and P35 and between P35 and P56. Most MR parameters in the gcc showed significant age-related changes except for T_1_ (**Fig. 2**). However, most MR parameters only showed significant differences between P14 and P35 but not between P35 and P56. Only FA and MD demonstrated significant differences across all age comparisons. In the cp, the difference in MOBP signal intensities between the later age groups (P35-P56) was significant while no significant difference was found between P14 and P35 (**Fig. 3**). Compared to the results from the gcc (**Fig. 2**), several MR parameters showed significant age-related changes, except T_1_, MTR, ihMTR, and RK. While T_2_, FA, MD, RD, and MK showed significant differences between P14 and P35, none of the MR parameters showed significant difference between P35 and P56. In the mCX, the MOBP signal intensities showed significant differences overall as well as between individual stages (**Fig. 4**). Among MR parameters, only T_2_, MK, and RK showed significant age-related difference and only between P14 and P35.

**Figure 2:**
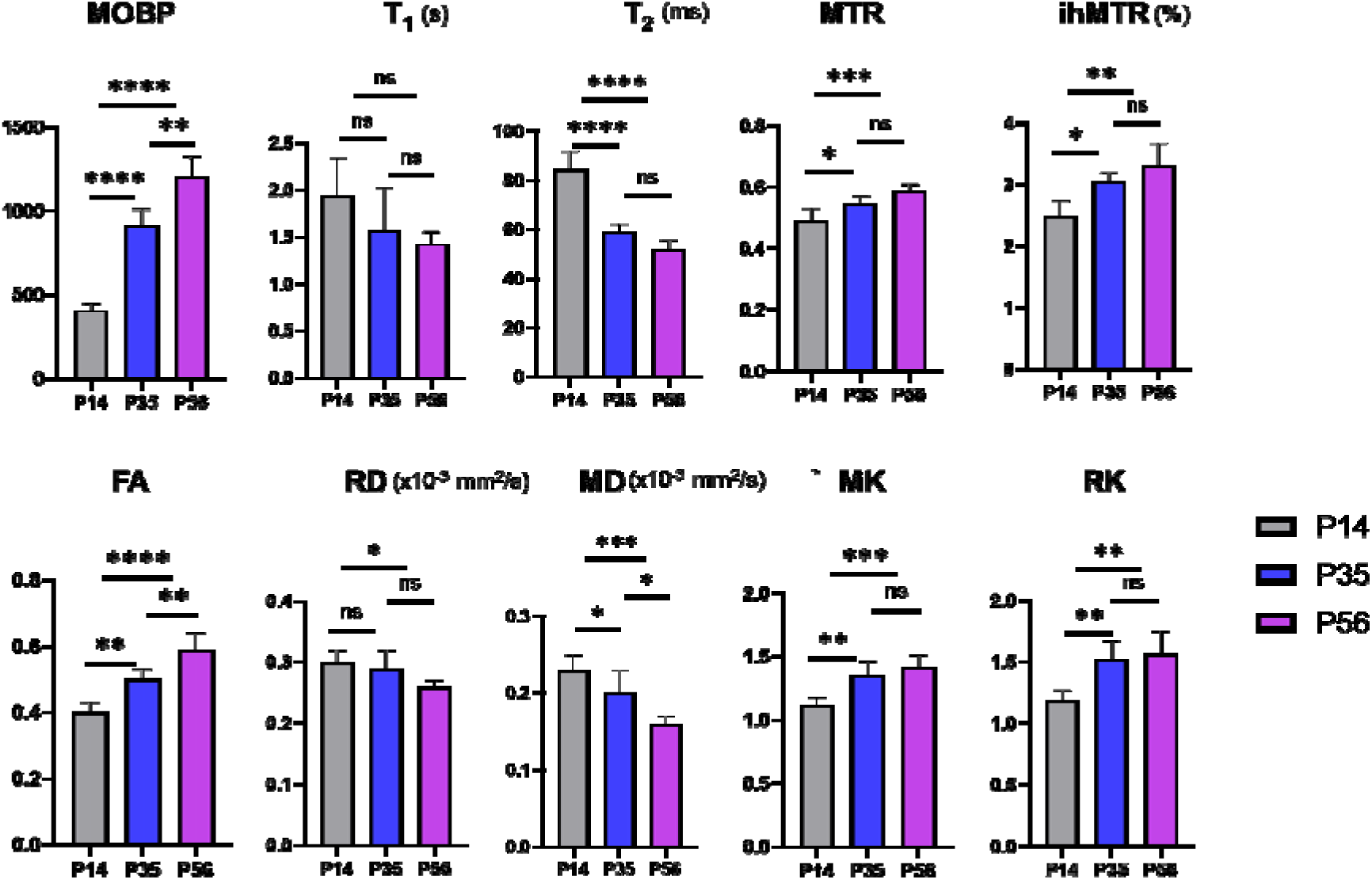
Mean ± standard deviation (SD) of the quantitative MRI parameters measured within each ROI for each developmental age group in the gcc from P14 to P56. The error bars indicate standard deviations. *, **, *** indicate p values less than 0.05, 0.01, and 0.001. The p values shown here are not corrected for multiple comparisons.

**Figure 3:**
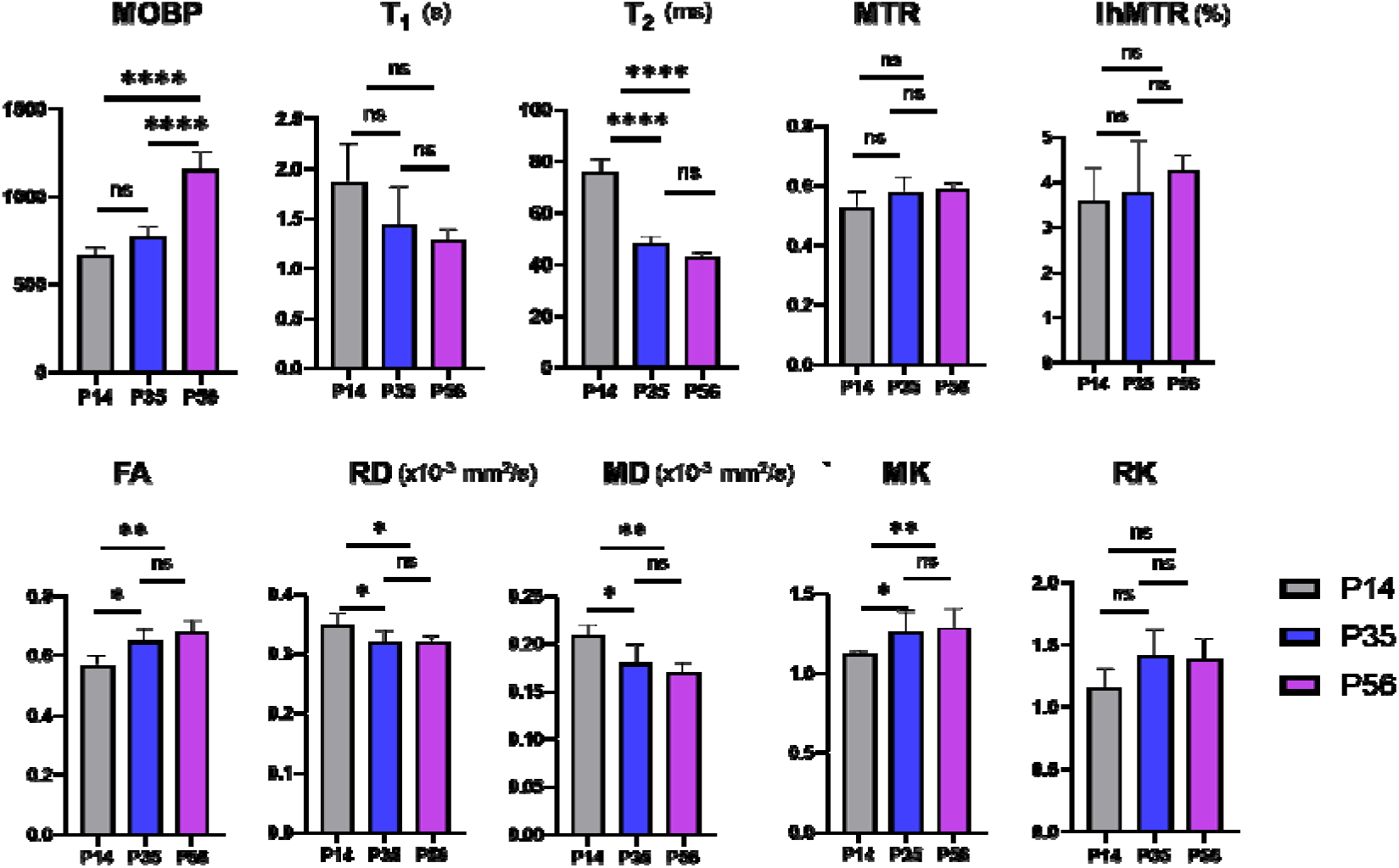
Mean ± standard deviation (SD) of the quantitative MRI parameters measured within each ROI for each developmental age group in the cp from P14 to P56. The error bars indicate standard deviations. *, **, *** indicate p values less than 0.05, 0.01, and 0.001. The p values shown here are not corrected for multiple comparisons.

**Figure 4:**
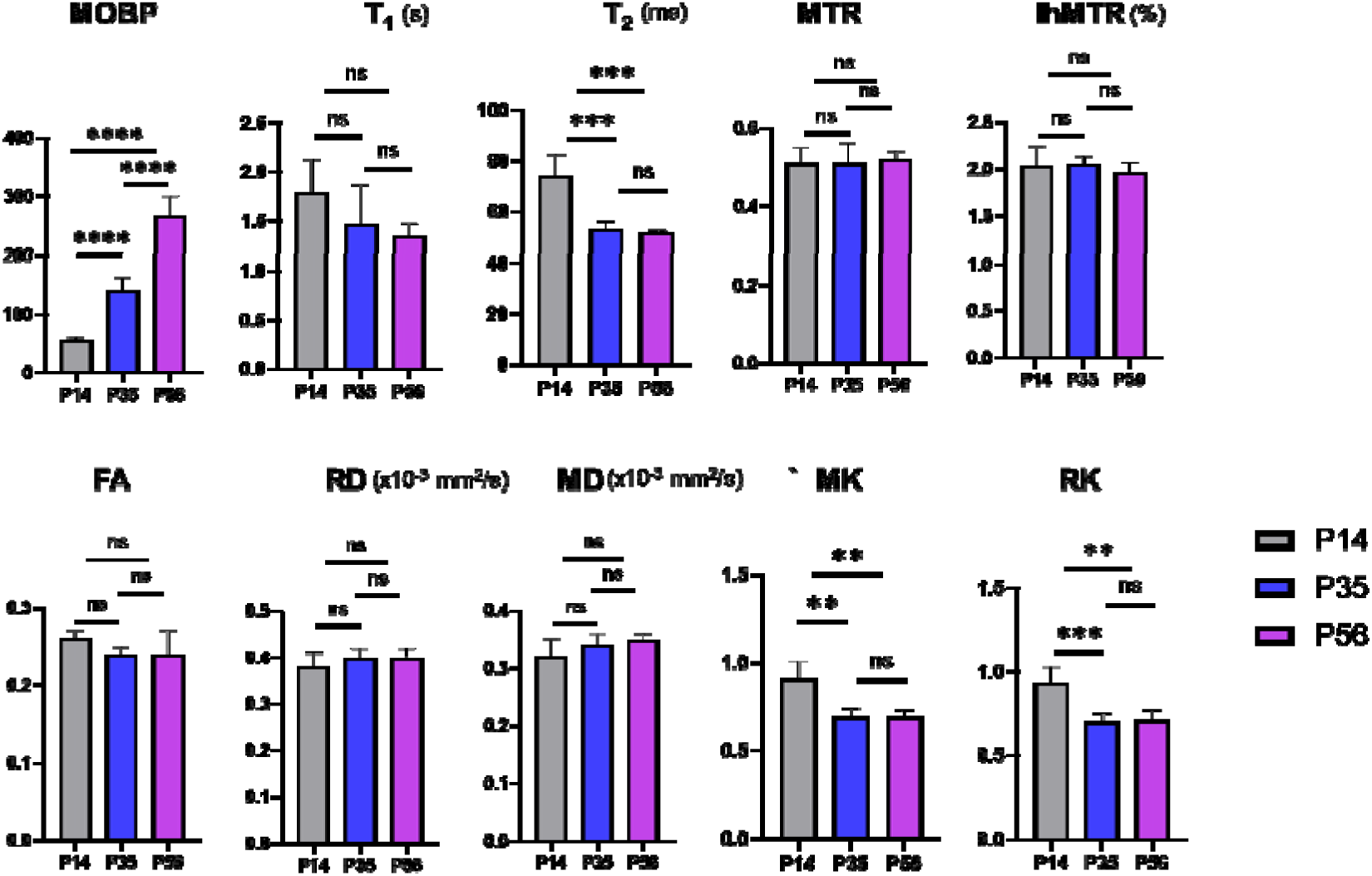
Mean ± standard deviation (SD) of the quantitative MRI parameters measured within each ROI for each developmental age group in the mCX from P14 to P56. The error bars indicate standard deviations. *, **, *** indicate p values less than 0.05, 0.01, and 0.001. The p values shown here are not corrected for multiple comparisons.

### 3.2 Correlations between MOBP and MR parameters

In the gcc, T_2_ had the strongest correlation with the MOBP signals ( =0.82, p<0.0001) (**Fig. 5**). FA and RD also showed strong correlations with the MOBP signals ( =0.78, p<0.0001 and =0.70, p=0.0002, respectively). T_1_, MTR, ihMTR, FA, and RK had moderate to strong correlations with the MOBP signals, but the correlations were not significant after corrections for multiple comparisons. In the cp, only T_2_ showed a significant correlation with the MOBP signals ( =0.53, p=0.003), whereas FA and RD showed moderate but insignificant correlations with the MOBP signals (**Fig. 6**). The other MR parameters showed weak and insignificant correlations with the MOBP signals. In the mCX, we only found a significant correlation between MOBP and T_2_ (**Fig. 7C**), with the rest of the MR parameters showing insignificant correlations (**Fig. 7**). In the cp and mCX, a nonlinear trend was observed between T_2_ and MOBP signals (**Fig. 6B** and **Fig. 7B**).

**Figure 5:**
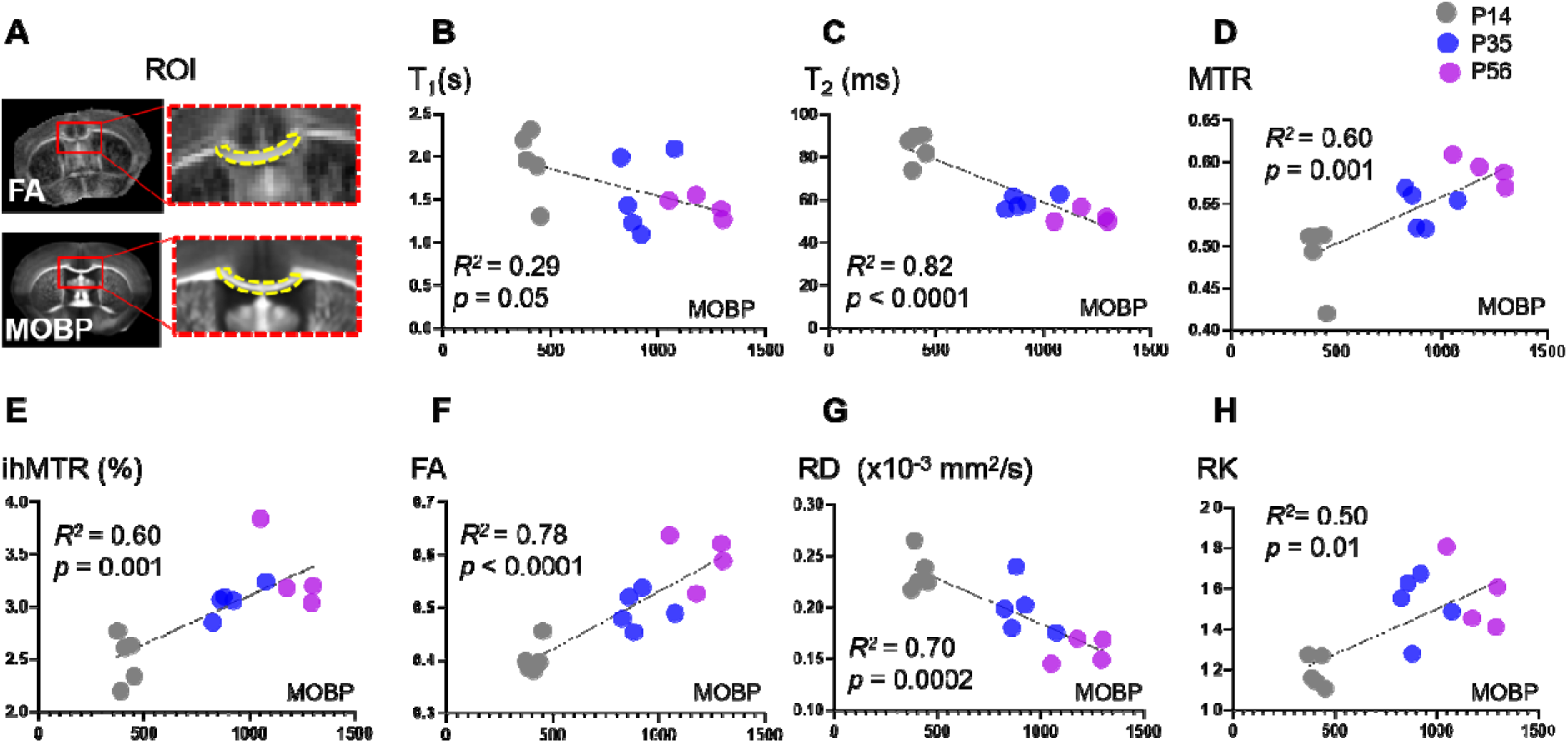
Correlation plots between individual MR parameters and the MOBP in the gcc. The defined ROI of gcc (pointed by dotted yellow outline) on the FA in the upper row and MOBP histology in the lower row in A. The p values shown here are not corrected for multiple comparisons.

**Figure 6:**
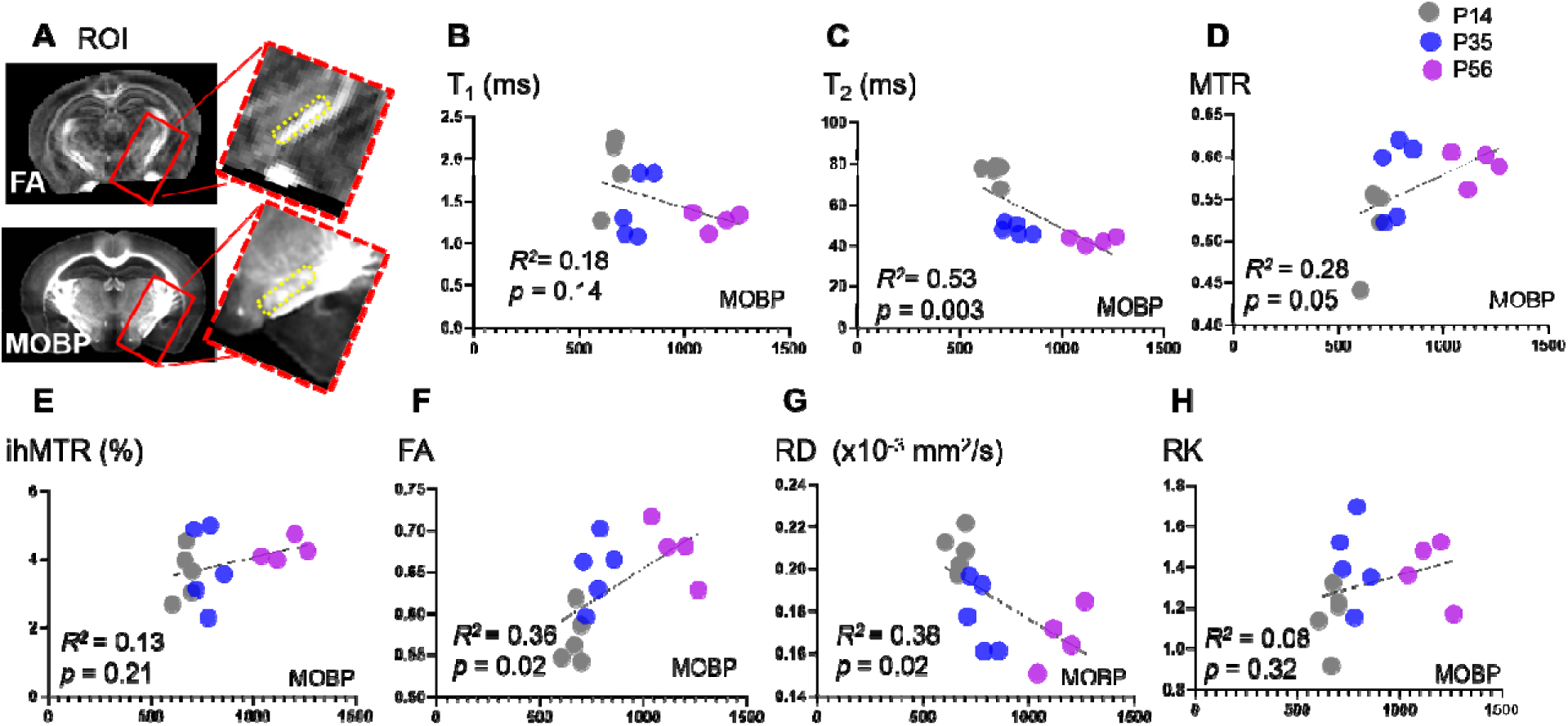
Correlation plots between individual MR parameters and the MOBP in the cp. The defined ROI of cp (pointed by dotted yellow outline) on the FA in the upper row and MOBP histology in the lower row in A. The p values shown here are not corrected for multiple comparisons.

**Figure 7:**
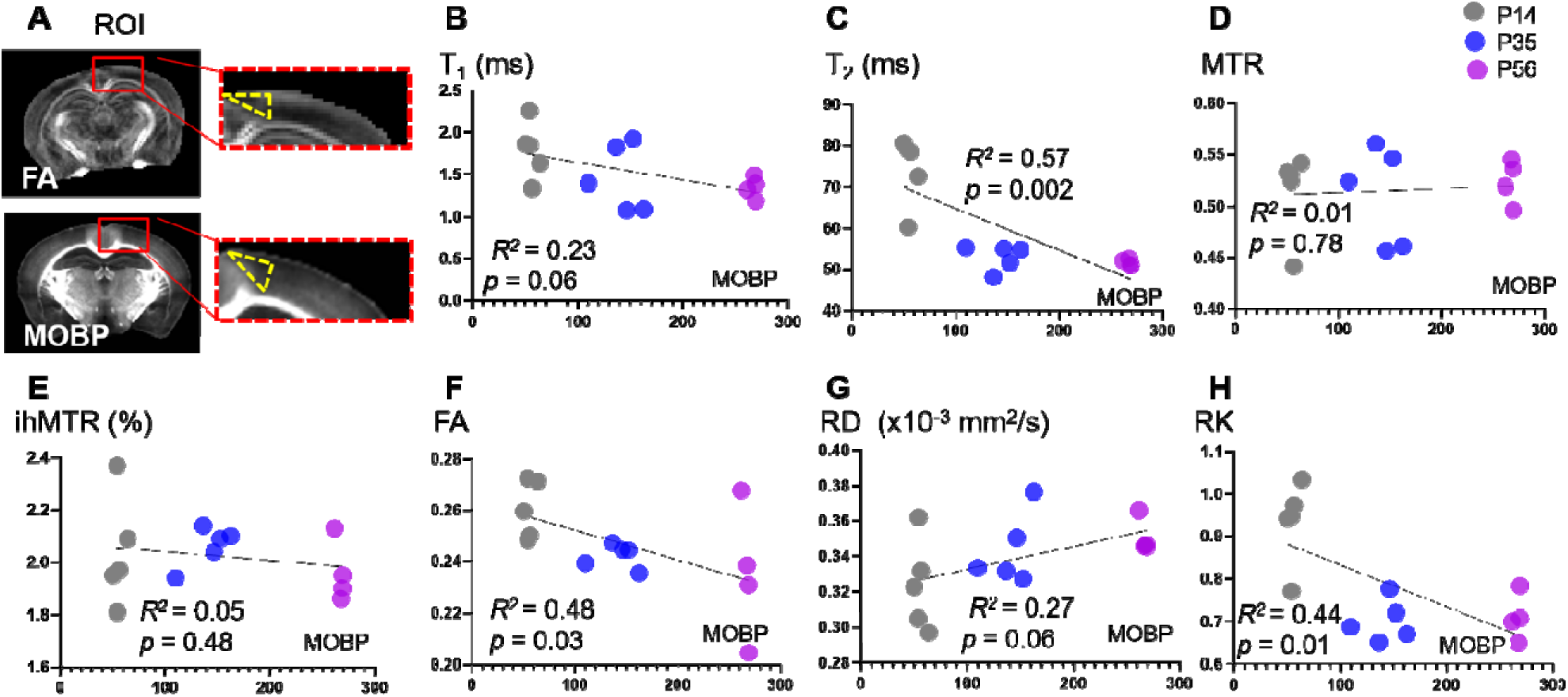
Correlation plots between individual MR parameters and MOBP in the mCX. The defined ROI of mCX (pointed by dotted yellow outline) on the FA in the upper row and MOBP histology in the lower row in A.

### 3.3 Contribution of individual MR parameter to myelin estimation

MRI parameters were included in the PLSR analysis based on their a priori relevance to complementary tissue properties associated with myelination, including relaxation, magnetization transfer, and diffusion-derived measures. Rather than selecting predictors based on visual differences in Figure 1 or their univariate association with MOBP, all quantitative MRI parameters that passed image-quality and data-completeness criteria were entered as candidate predictors. PLSR was used because it is well suited for modeling correlated predictors by projecting them onto a smaller number of latent components that maximize covariance with the response variable. This approach minimized subjective parameter selection while allowing the relative contribution of each MRI measurement to MOBP variation to be evaluated within the multivariate model. For the gcc, T_2_ has the highest VIP score (**Fig. 8A**, with one latent factor in PLSR). When all seven MR parameters were included, up to 90% of the variance in MOBP signals from the gcc was explained with two latent factors in PLSR (the blue curve in **Fig. 8B**). Similar performance was achieved with only T_2_, FA, and RD with two latent factors (the green curve in **Fig. 8B**). With T_2_ only or T_2_ plus MTR, the amount of variance in MOBP signals from the gcc that can be explained was slightly reduced (the orange curve in **Fig. 8B**). The myelin estimator based on the PLSR result from gcc (**Tables 2 & 3,** all seven MR parameters, one latent factor in PLSR) showed strong correlation with MOBP signals (R^2^ = 0.93 and p <0.0001) (**Fig. 8C**). In the cp, T_2_ also had the highest VIP score (**Fig. 8D**, with two latent factors in PLSR). Three latent components in PLSR were needed to explain approximately 60% variance of the MOBP signals (**Fig. 8E**), with minute improvement by adding additional latent components. The myelin estimator based on the PLSR result (**Table 2**) showed strong correlation with MOBP signals in the cp (*R*^2^= 0.64 and p = 0.0006) (**Fig. 8F**). In the motor cortex (mCX), T_2_ again had the highest VIP score as well (**Fig. 8G**), and three latent components were needed to explain approximately 70% variance of the MOBP signals (**Fig. 8H**). The myelin estimator based on the PLSR result showed strong correlation with MOBP signals in the mCX (*R*^2^=0.74 and p < 0.0001) (**Fig. 8I**). For all three structures, the PLSR generated myelin estimators had stronger correlations with the corresponding MOBP signals than any single MR parameters.

**Fig. 8:**
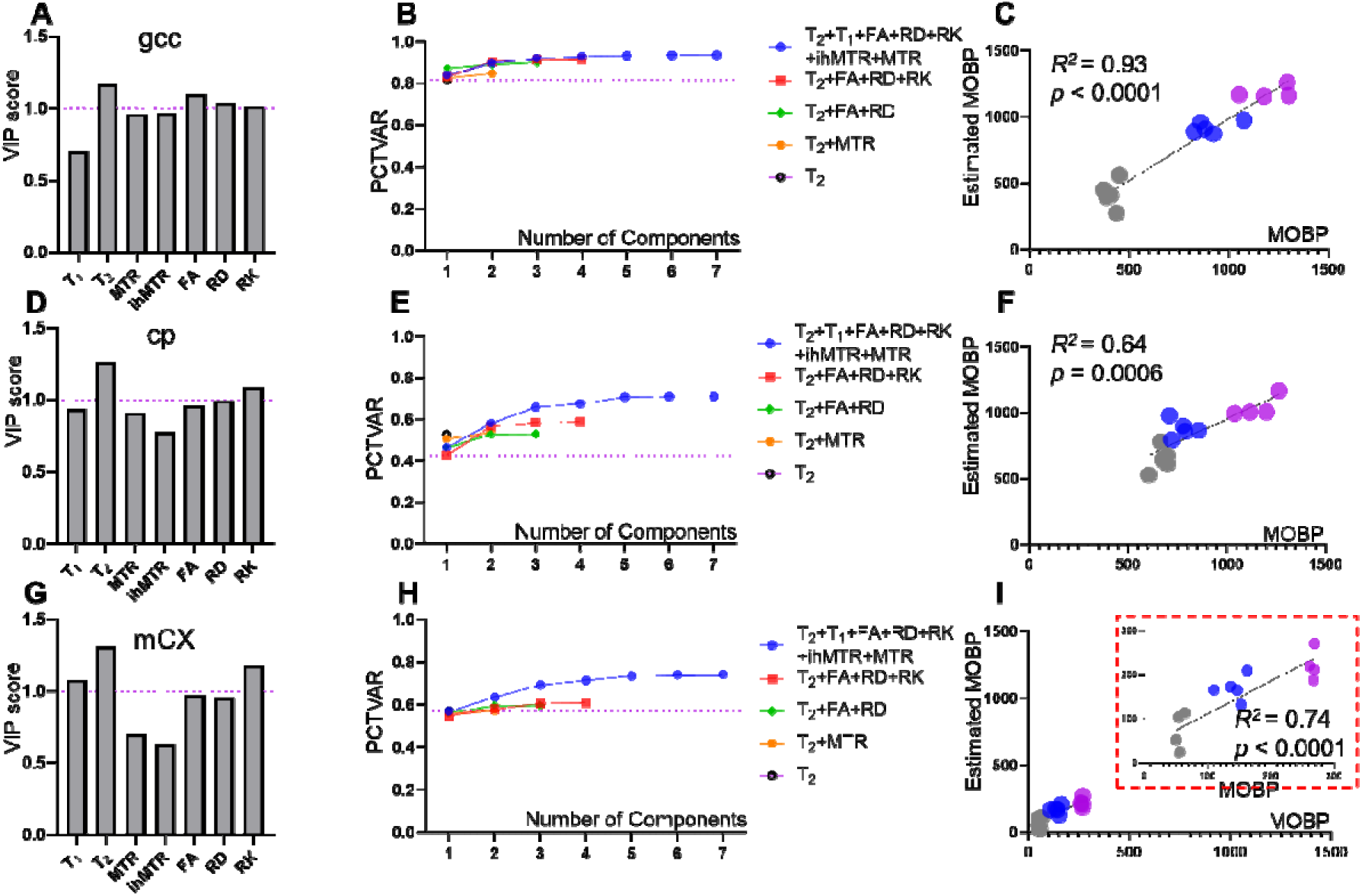
PLSR analysis of MRI markers predicting MOBP across different brain regions. The PLSR analysis provided the VIP scores of individual MR parameters, the percentage of variance in MOBP signal intensities that can be explained by the latent components for the gcc (top row), cp (middle row), and mCX (bottom row). The correlations between the estimated MOBP signals based on PLSR and actual MBOP signals were shown in the right column.

**Table 2.** PLSR coefficients in different ROIs using 7 markers. (LC: latent component)

| ROI | No. of LC | Constant | T<sub>1</sub> | T<sub>2</sub> | MTR | ihMTR | FA | RD | RK |
| --- | --- | --- | --- | --- | --- | --- | --- | --- | --- |
| gcc | 1 | 2177 | -0.32 | -20.5 | 0.06 | 0.48 | 0.11 | -0.05 | 0.23 |
| cp | 2 | 1717 | -198.9 | -8.7 | -4.07 | 138.69 | -30.62 | 7.46 | -442.65 |
| mCX | 2 | 417.07 | -73.79 | -4.95 | 55.25 | 51.45 | -14.73 | -4.81 | 17.84 |

**Table 3.** PLSR coefficients in different ROIs using 7 markers after standardization.

| ROI | No. of LC | Constant | T<sub>1</sub> | T<sub>2</sub> | MTR | ihMTR | FA | RD | RK |
| --- | --- | --- | --- | --- | --- | --- | --- | --- | --- |
| gcc | 1 | 0 | -0.11 | -0.18 | 0.15 | 0.15 | 0.17 | -0.16 | 0.14 |
| cp | 2 | 0 | -0.34 | -0.42 | 0.28 | 0.28 | 0.07 | -0.13 | -0.51 |
| mCX | 2 | 0 | -0.52 | -0.38 | 0.29 | 0.22 | -0.19 | 0.02 | -0.06 |

To test generalizability, the performance of the myelin estimator derived from PLSR based on data from the gcc (**Fig. 9A-C**) was evaluated using the data from the isthmus of the corpus callosum (icc). With all 7 parameters and one latent component, we found a strong correlation with *R* ^2^of 0.94 (**Fig. 9A**). When we reduced the number of MR parameters to T_2_ and RD only (**Fig. 9D**), selected based on the VIP scores, we still had strong correlations (*R*^2^of 0.97, 0.85, respectively). The PLSR derived linear myelin estimators are MOBP = 1970.93 -23.18*T_2_ + 713.96*RD. In the cp, prediction using the myelin estimator based on the gcc data only showed a moderate correlation with the MOBP signal intensities (*R*^2^=0.57 vs. 0.53, **Fig. 9B, and E**), with little added benefit from more parameters. In mCX, correlations were modest when using all seven parameters (*R*^2^=0.41) but improved when restricted to two parameters (*R*^2^=0.58) (**Fig. 9C, and F**).

**Fig. 9:**
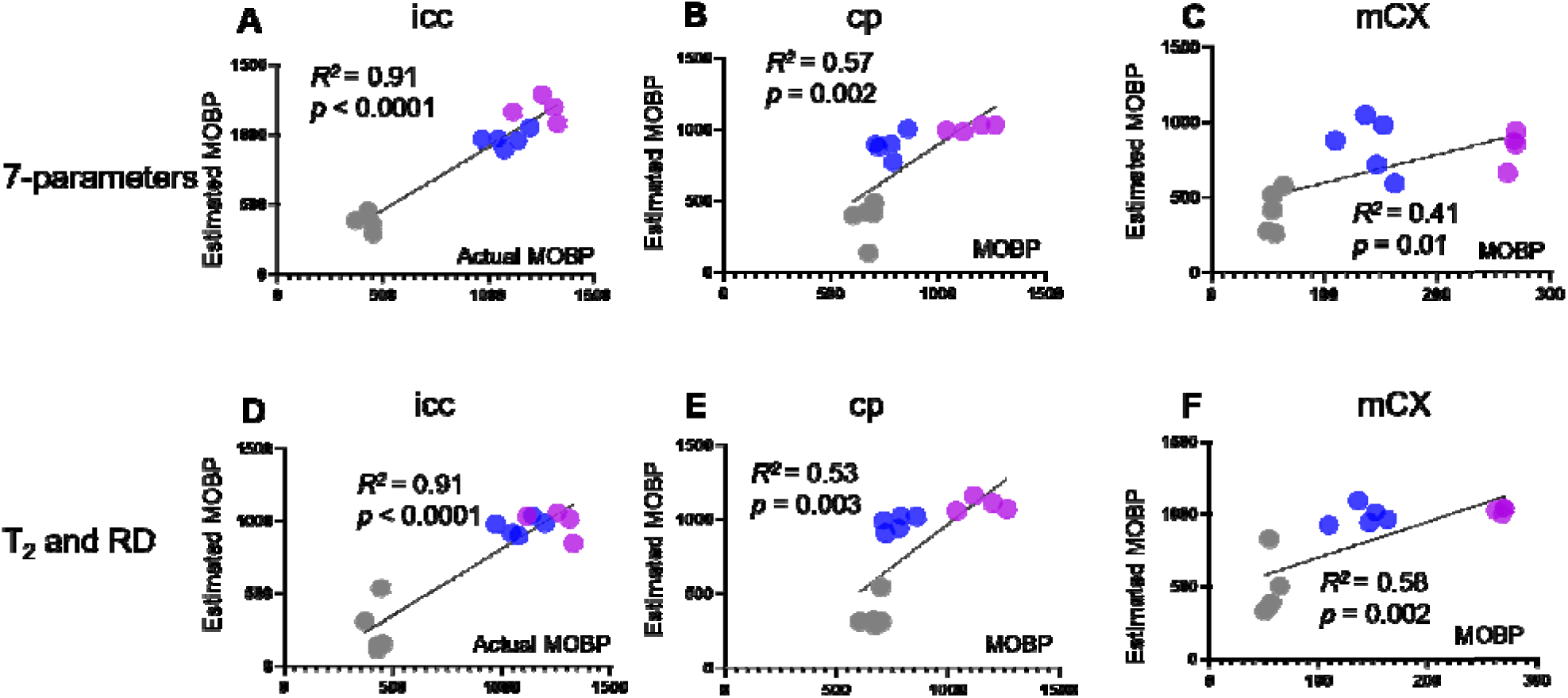
Correlation plots between actual and estimated MOBP signals in the icc, cp, and mCX using the PLSR results in **Table 3**.

## 4. Discussion

### 4.1. The use of MOBP signal intensity as the reference tissue myelin content

In this study, we used MOBP signal intensity from STPT data as the histological reference to evaluate the performance of mpMRI for characterizing developmental myelination. Compared with classical myelin markers such as MBP and PLP, MOBP is expressed specifically in the CNS and appears approximately 2–3 days later than MBP during postnatal development (61, 62).

Based on semi-quantitative Western blot measurements, MOBP accounts for approximately 5– 10% of the level of MBP in CNS myelin (61, 63, 64), although the precise relationship between MOBP and MBP across developmental stages is not known. This difference in developmental timing is particularly relevant at P14, when active myelination is still ongoing. MOBP expression at this early time point may therefore underrepresent total myelin relative to earlier-expressed markers such as MBP. Consequently, the lower MOBP signal at P14 could increase the apparent dynamic range between P14 and later developmental stages and thereby contribute to the magnitude of MRI–MOBP associations observed across ages. Thus, developmental increases in MOBP should not be interpreted solely as increases in total myelin content, but may also reflect ongoing oligodendrocyte and myelin maturation.

Despite this limitation, the MOBP-eGFP mouse provides an important advantage for evaluating spatial patterns of myelination throughout the brain. Previous studies have used MOBP-eGFP mice to investigate activity-dependent myelin plasticity (65), demyelination and remyelination in the cuprizone model (66, 67), and the genetic regulation of oligodendrocyte maturation (68). Whole-brain STPT imaging of MOBP-eGFP brains also overcomes several practical limitations of conventional light microscopy, including limited brain coverage associated with labor-intensive sectioning and staining, tissue deformation or damage, and variability introduced by histological staining. The spatially continuous STPT dataset was particularly advantageous for the present study because it enabled consistent comparisons of MRI–MOBP relationships across multiple white- and gray-matter regions.

Nevertheless, MOBP represents one component of myelin biology rather than a developmentally invariant measure of total myelin content. Accordingly, the strong association between T_2_ and MOBP observed in this study should be interpreted as sensitivity of T_2_ to tissue changes accompanying MOBP-associated developmental myelin maturation, rather than evidence that T_2_ is a direct or specific measure of total myelin. More quantitative assessment of myelin ultrastructure will require complementary measurements, such as electron microscopy (EM)-derived myelin thickness and g-ratio. However, EM is typically restricted to relatively small tissue regions, making comprehensive comparisons across brain regions and developmental stages challenging. Future studies integrating MOBP-STPT with additional myelin markers and ultrastructural measurements will therefore be important for determining whether the MRI–MOBP relationships identified here generalize to other measures of myelin content and maturation.

### 4.2 Region-specific correlations

Given the complexity of the biological microstructures and many nuisance factors varying across regions of the brain, it is important to minimize confounding factors and focusing on correlations between MRI and histological measurements within the same anatomical structures (41–43). To further illustrate this point, our analysis on data from gcc, cp, mCX, showed distinct relationship between MR parameters and MOBP signals (**Fig. 8**).

An important finding of this study is that the predictive performance of the PLSR model depended on the anatomical region to which it was applied. A model trained in the gcc showed relatively preserved performance when applied to the icc; however, because both regions are callosal white matter with broadly similar tissue organization, this result is more appropriately interpreted as within-callosal transferability rather than evidence of broad anatomical generalizability. In contrast, application of the gcc-trained model to the cp and particularly the mCX resulted in substantially lower predictive performance, demonstrating that an MRI–MOBP relationship learned in one white-matter region cannot be assumed to generalize across anatomically and microstructurally distinct regions.

This regional dependence is expected given the nonspecific and overlapping biological sensitivities of the MRI parameters included in the model. As myelin and other tissue microstructures (e.g. axons) are tightly coupled, it is difficult to completely remove potential confounding factors while preserving changes in myelin. For example, parallel to myelination in the mouse brain, there are processes (e.g. axonal pruning and synaptic remodeling (69), gliogenesis (70), iron accumulation (71), and water content shifts overlap with myelination (72)). All these processes can also affect the measured MR parameters and lead to potential over-estimation. Consequently, the relative PLSR weights derived in the gcc likely reflect both myelination and the specific microstructural environment of callosal white matter. The reduced cross-region performance therefore highlights an important limitation of region-specific multivariate models and suggests that broader myelin mapping will require models trained across anatomically diverse white- and gray-matter regions or explicit consideration of regional tissue characteristics.

Another consideration is the collinearity among MR parameters. MRI often suffers from multi-collinearity because most of them are not specific to a single cellular compartment (e.g. myelin). Instead, they are influenced by overlapping biophysical and microstructural factors. For example, relaxometry (T_1_, T_2_) is affected by water content, myelin, and iron(44, 73–75). Magnetization transfer reflects macromolecular content but is also influenced by water relaxation times(76–79). The use of PLSR can accommodate the collinearity but is limited by its linear assumption. For example, the relationship between T_2_ and MOBP signals in the cp and mCX showed a nonlinear trend with age compared to the gcc. This may explain the lower performance of PLSR-based myelin estimators for the cp and mCX than the gcc.

### 4.3 T_2_ as a main driver for myelin estimation

Across all three regions—the gcc, cp, and mCX—PLSR consistently identified T_2_ as the dominant MR parameter (highest VIP) explaining most of variances in MOBP signals. This is not surprising given the strong correlations between T_2_ and MOBP signals consistent with histological validation studies demonstrating robust correlations between T_2_-derived indices and myelin content across white and gray matter (15, 80). Previous reports including a systemic review paper(43) align well with our observation that T_2_ remains robust across tissue classes to reflect the myelin level in human brain (12) and in mouse models (80). Several factors may explain the strong association between T_2_ and MOBP during postnatal development. Myelination alters both the fractional distribution and exchange dynamics of short- and intermediate-T_2_ water pools (81–83). Increased myelin compaction reduces interlamellar water and restricts water mobility, leading to predictable shortening of T_2_ (84, 85). Increasing number of oligodendrocytes, which produce MOBP and are rich in iron may also shorten T_2_. All these factors potentially contribute to the strong contribution by T_2_ to myelin estimation.

However, including additional MR parameters can improve our ability to estimate myelin content across WM and GM. The improvements were clear in the cp and mCX (**Fig. 8**), with noticeable gap between the amount of variance in MOBP signals that can be explained by T_2_ alone or multiple MR parameters. This is also supported by the relative VIP scores of other MR parameters (**Fig. 8D & G**). The results suggest that diffusion-based metrics may provide complementary information for myelin estimation, which is likely due to strong coupling between axon and myelin and the sensitivity of diffusion MRI to the restrictive effects from axon membrane and myelin sheath as well as axonal density and orientation dispersion (86–88). In the cortex, T_1_ also had VIP scores higher than 1, indicating strong contribution to myelin estimation. This aligns with reports that using the ratio between T_1_-weighted and T_2_-weighted MRI to map cortical myelination(89–91).

### 4.4 Limitations

Several limitations of this study should be noted. First, the current study was performed on post-mortem brain specimens. Death and chemical fixation alter absolute MR parameters (92, 93), including shortened T_1_ and T_2_ and lower diffusivity. Additionally, postnatal developing mouse brains lack a neuroinflammation component, which can alter tissue microstructures and related MR parameters. The human brain and mouse brain whit matter also have distinct microstructures (e.g. axon diameter and degree of myelination) (94, 95).

An important limitation of this study is that all MRI measurements were acquired ex vivo following PFA fixation. Fixation alters tissue water compartmentalization and molecular mobility and consequently affects quantitative relaxation and diffusion measurements. Therefore, the absolute MRI values obtained here cannot be directly compared with in vivo measurements. Importantly, fixation may affect individual MRI contrasts differently; thus, the relative contributions of MRI parameters identified by PLSR, including the predominance of T2 and the corresponding VIP rankings, should not be assumed to remain unchanged in living tissue. The present results should instead be interpreted as MRI–MOBP relationships established under controlled fixed-tissue conditions. Their value for in vivo myelin imaging will require direct validation using longitudinal in vivo MRI followed by spatially matched ex vivo MRI and histological measurements in the same animals.

Second, the present analysis did not include some advanced myelin markers such as the myelin water fraction (MWF) (96, 97) primarily for technical reasons. Despite the high specificity of MWF to myelin (12, 78), accurate estimation of MWF remains challenging. Multi-echo acquisitions required for MWF mapping are time-consuming and highly sensitive to signal-to-noise ratio and B_0_/B_1_ inhomogeneity, often limiting spatial resolution or whole-brain coverage (98, 99). With limited availability of the scan times, only 25 echoes with 7ms spacing was possible but insufficient to perform MWF analysis (with shorter T_2_ in fixed tissues). The observed strong contribution of T_2_ to myelin estimation and previous reports on MWF under neuroinflammation suggest that including MWF in future studies will be beneficial.

## Conclusion

This study demonstrates that multi-parametric MRI can characterize developmental myelination in the ex vivo mouse brain. T_2_ provided the strongest and most consistent contribution to explaining MOBP variance across white and gray matter, potentially reflecting changes in myelin-associated water compartmentalization during myelin maturation. Complementary contributions from other MRI parameters further support a multi-parametric approach for capturing the complex tissue changes accompanying myelin development.

## Acknowledgement

This study was supported by the National Institute of Health R01NS102904, R01HD074593, U24NS135568. MR imaging was performed at the Preclinical Imaging Center supported by 1S10OD018337-01 and Center of Advanced Imaging Innovation and Research (CAI2R, www.cai2r.net), a Biomedical Technology Resource Center supported by NIBIB with the award P41 EB017183.

