## Supplemental figures 1-3 for "Multi-parametric ex vivo magnetic resonance imaging for myelination in the developing mouse brain"

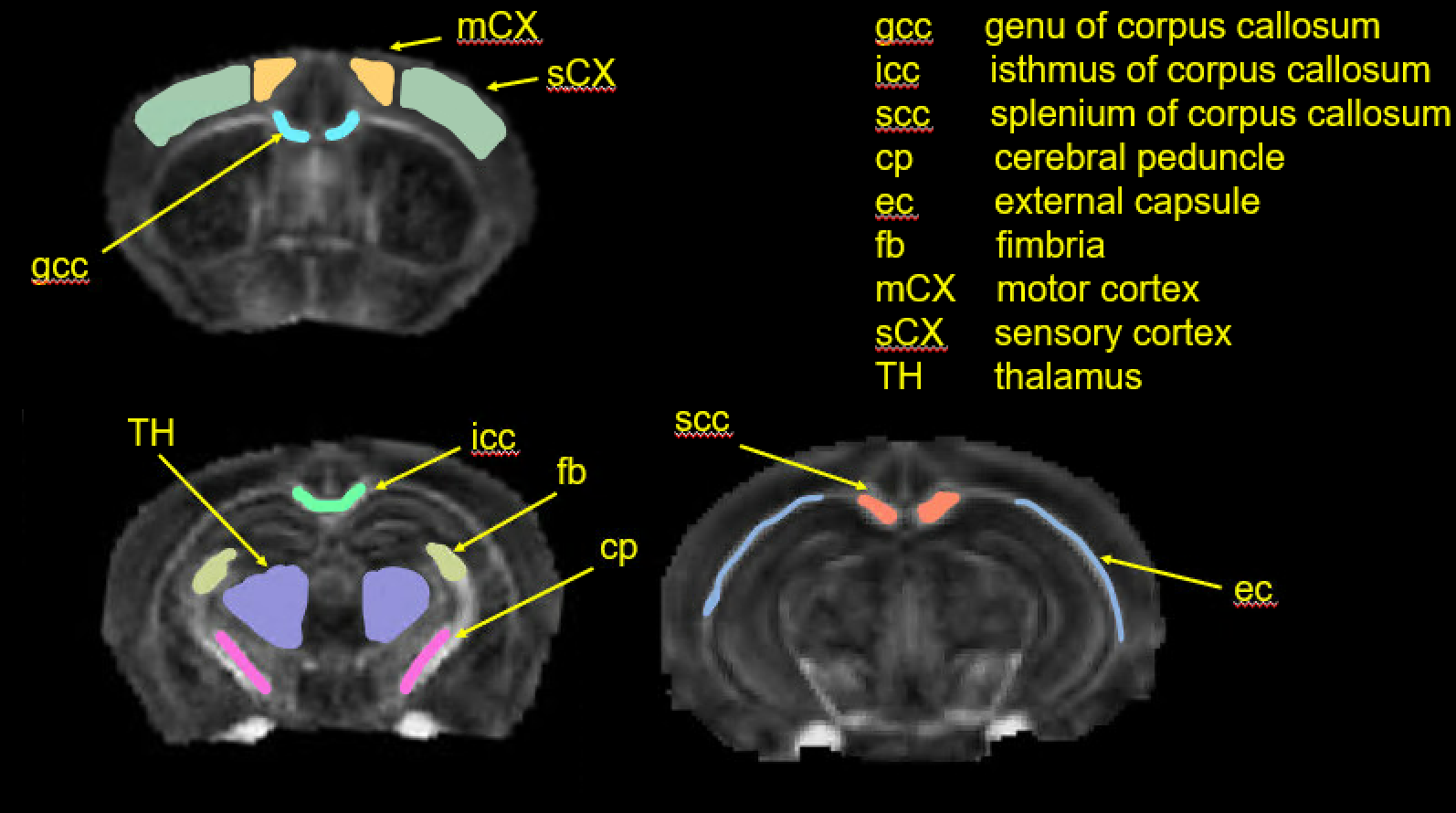

**Fig. S1:** The representative ROIs were defined manually in the FA image of a P56 mouse brain. The total ROIs consists of 6 WM and 3 GM ROIs.

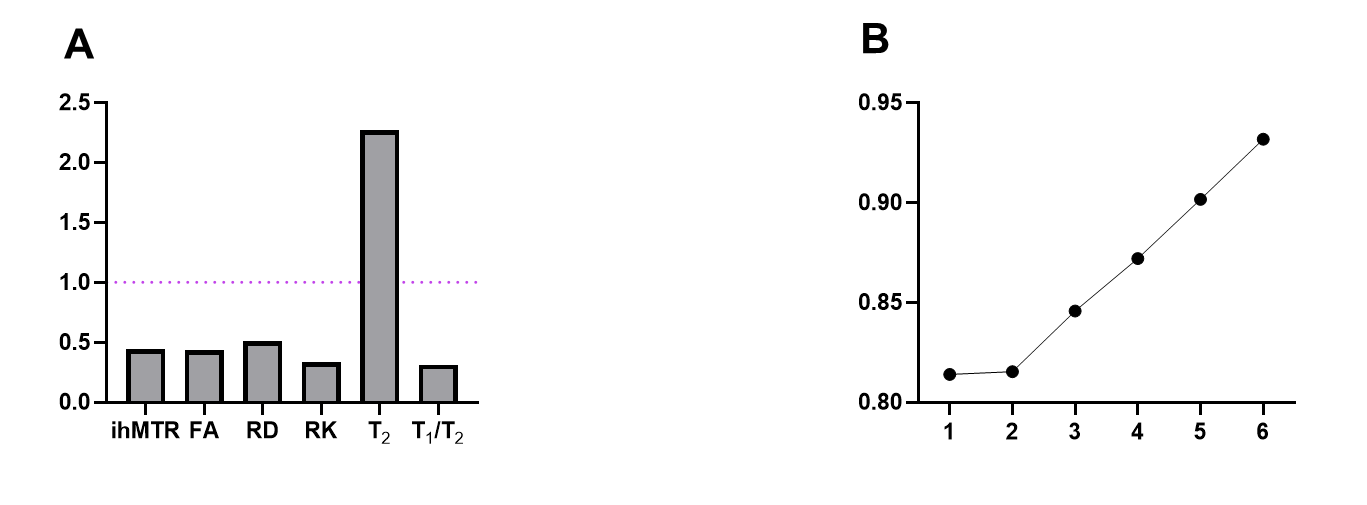

**Fig. S3**: Estimation of myelin based on multiple MR parameters in the gcc. **A-B:** The variable importance in projection (VIP) scores of each MR parameter and the percentage of variance (PCTVAR) of the MBP signals in the gcc that can be explained by latent components.

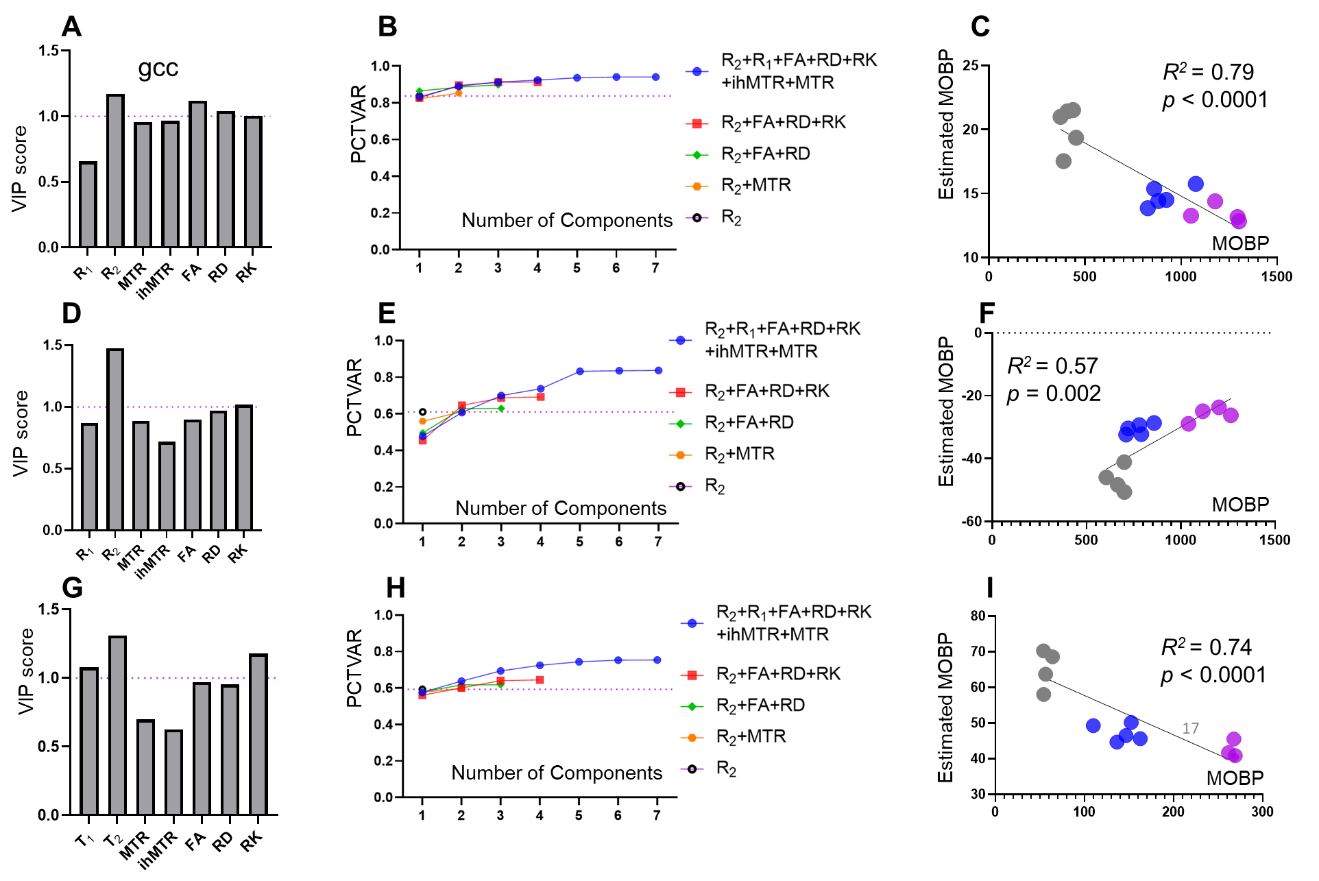

**Fig. S4: PLSR analysis of MRI markers predicting MOBP across different brain regions.**
The PLSR analysis provided the VIP scores of individual MR parameters, the percentage of variance in MOBP signal intensities that can be explained by the latent components for the gcc (top row), cp (middle row), and mCX (bottom row). The correlations between the estimated MOBP signals based on PLSR and actual MBOP signals were shown in the right column.

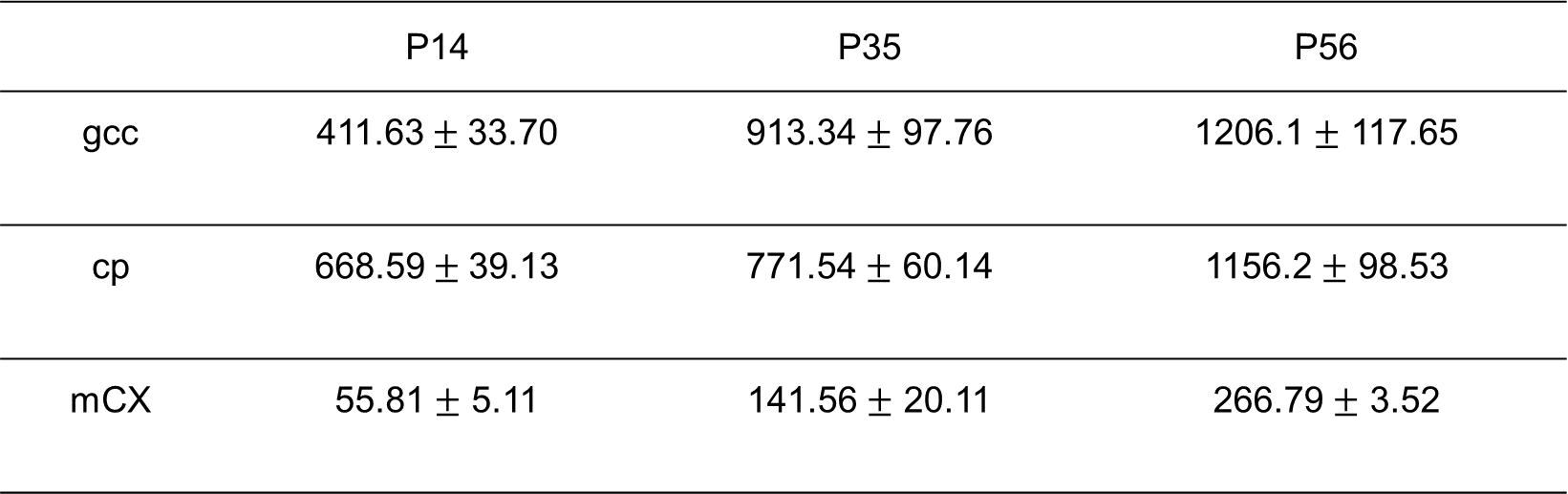

**Table1:** The MOBP signal intensity at different age groups of MOBP-eGFP mouse in STPT

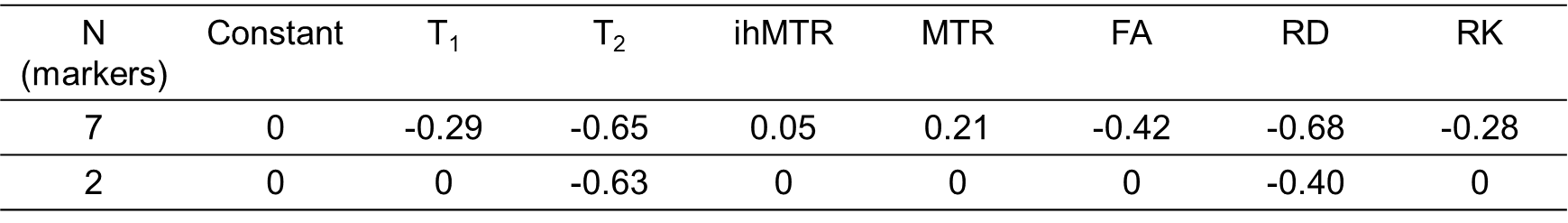

**Table2:** The linear coefficients in markers with different combinations: 7, 2 markers.
